# Short-term Topiramate treatment prevents radiation-induced cytotoxic edema in preclinical models of breast-cancer brain metastasis

**DOI:** 10.1101/2023.02.14.528559

**Authors:** Maria J. Contreras-Zárate, Karen LF. Alvarez-Eraso, Jenny A. Jaramillo-Gómez, Zachary Littrell, Niki Tsuji, David R. Ormond, Sana D Karam, Peter Kabos, Diana M. Cittelly

## Abstract

**Background:** Brain edema is a common complication of brain metastases (BM) and associated treatment. The extent to which cytotoxic edema, the first step in the sequence that leads to ionic edema, vasogenic edema and brain swelling, contributes to radiation-induced brain edema during BM remains unknown. This study aimed to determine whether radiation-associated treatment of BM induces cytotoxic edema and the consequences of blocking the edema in pre-clinical models of breast cancer brain metastases (BCBM).

**Methods:** Using *in vitro* and *in vivo* models, we measured astrocytic swelling, trans-electric resistance (TEER) and aquaporin 4 (AQP4) expression following radiation. Genetic and pharmacological inhibition of AQP4 in astrocytes and cancer cells was used to assess the role of AQP4 in astrocytic swelling and brain water intake. An anti-epileptic drug that blocks AQP4 function (topiramate) was used to prevent cytotoxic edema in models of BM.

**Results:** Radiation-induced astrocytic swelling and transient upregulation of AQP4 within the first 24 hours following radiation. Topiramate decreased radiation-induced astrocytic swelling, loss of TEER in astrocytes *in vitro*, and acute short term treatment (but not continuous administration), prevented radiation-induced increase in brain water content without pro-tumorigenic effects in multiple pre-clinical models of BCBM. AQP4 was expressed in clinical BM and breast cancer cell lines, but AQP4 targeting had limited direct pro-tumorigenic or radioprotective effects in cancer cells that could impact its clinical translation.

**Conclusions:** Patients with BM could find additional benefits from acute and temporary preventive treatment of radiation-induced cytotoxic edema using anti-epileptic drugs able to block AQP4 function.

**Key points:**

- Radiation induces cytotoxic edema via acute dysregulation of AQP4 in astrocytes in preclinical models of BM.
- Pharmacologic blockage of AQP4 function prevents water intake, astrocytic swelling and restores TEER *in vitro.*
- Pre-treatment with single-dose Topiramate prevents brain radiation-induced brain edema without direct tumor effects in pre-clinical models of BCBM.

**IMPORTANCE OF THE STUDY:** In this study we describe a novel role for astrocytic swelling and cytotoxic edema in the progression of radiation-induced brain edema during BM treatment. While radiation-induced edema has been fully attributed to the disruption of the blood-brain barrier (BBB) and ensuing vasogenic effects, our results suggest that cytotoxic edema affecting astrocytes in the acute setting plays an important role in the progression of brain edema during BM standard of care. Current standard of care for brain edema involves pre-treatment with steroids and the use of bevacizumab only after clinically significant edema develops. Both interventions are presumed to target vasogenic edema. This study suggests that patients with BM could find additional benefits from acute and temporary preventive treatment of radiation-induced cytotoxic edema using an already FDA-approved anti-epileptic drug. Such early prevention strategy can be easily clinically implemented with the goal of minimizing treatment-related toxicities.

## INTRODUCTION

Brain metastases (BM) represent the most common intracranial malignancy in adults, and breast cancer is a frequent primary site leading to BM. Standard of care for BM includes stereotactic radiosurgery alone or in combination with surgery, systemic chemotherapy or targeted therapies^1^. A common complication of BM and their treatments is brain edema, a pathological state in which brain volume increases due to abnormal accumulation of fluid and ions within the cerebral parenchyma^2^. Unlike edema in other organs, brain edema can be lethal or significantly impair life quality due to intracranial hypertension.

Brain edema associated with BM and radiation is attributed mainly to vasogenic edema, a disruption of the BBB resulting in the extravasation of serum components and entry of fluids to the interstitial space from the bloodstream^3^. However, a first step in the sequence that leads to ionic edema, vasogenic edema, and ultimate brain swelling and necrosis^4,5^, is cytotoxic edema, a premorbid cellular process, whereby extracellular Na^+^ and other cations enter into neurons and astrocytes and accumulate intracellularly, resulting in osmotic expansion of the cells and leading to necrotic cell death^3,4,6^. While cytotoxic edema has been described in spinal cord injury, traumatic brain injury and stroke, the extent to which cytotoxic edema contributes to the pathophysiology of BM and their treatments remains unknown.

Cytotoxic edema is characterized by astrocytic swelling and dysregulation of AQP4 levels, the main bilateral water channel in the central nervous system (CNS)^7–10^. AQP4 is expressed abundantly in astrocytes and plays different roles in development and resolution of CNS edema, with water flow through AQP4 driving the development of cytotoxic edema in the early post-injury stage but later clearing vasogenic edema^11,12^. Therefore, reversible inhibition of AQP4 function during the acute phase of CNS trauma has been proposed as a viable strategy to prevent CNS edema^13,14^. We have shown that radiation induces AQP4 dysregulation in immortalized astrocytes *in vitro* and that combination of radiation with cytotoxic drugs (i.e T-DM1 used to treat Her2+ BM)^15^ further increases AQP4 dysregulation and astrocytic cell death, suggesting AQP4 and cytotoxic edema play a role in radiation-induced edema in BM. Herein, we demonstrate that radiation induces AQP4 dysregulation and cytotoxic edema in human primary astrocytes *in vitro* and *in vivo*, and show that an FDA-approved anti-epileptic drug Topimarate, reported blocking AQP4 function^16^ can be used to prevent radiation-induced cytotoxic edema in pre-clinical models of BCBM.

## MATERIALS AND METHODS

### Cells

HuAST (primary human astrocytes from ScienceCell #1800-10), HAL (immortalized human astrocytes from Dr. Patricia S. Steeg, NCI, US), THV (immortalized human astrocytes, from Dr. Paul B. Fisher). Human breast cancer cells were: BT474 (from Dr. Dihua Yu, MD Anderson Cancer Center, TX, USA); MCF-7 (RRID: CVCL_0031), JmT1BR3 and 231BR (from Dr. Patricia Steeg, NIH, USA); and F2-7cells^17^. Murine mammary gland tumor cells were: 4T1BR5 (derived from 4T1 from Dr. Suyun Huang, MD Anderson Cancer Center, TX, USA) and E0771BR (derived from E0771 RRID: CVCL_GR23). Details in supplemental methods.

### Human-derived BM

De-identified paraffin-embedded archived BM samples from female breast cancer patients were obtained from consenting donors under approved IRB protocol 20-1669 at the University of Colorado.

### Drugs

Topiramate (TPM, Selleckchem S7998) was used *in vitro* at 10-100 μM and *in vivo* at 50 mg/Kg/day in 10% DMSO. N-(1,3,4-thiadiazol-2-yl) pyridine-3-carboxamide dihydrochloride (TGN-020, Tocris Cat No 5425) was used *in vitro* at 10-100 μM. T-DM1 (Genetech) was used *in vitro* at 1 μg/ml.

### Cell proliferation

1,000-3,000 cells were plated in 96 well plates, pretreated for 2 hours before radiation when indicated, and imaged over time using Live Cell Incucyte Imaging (Essen Bioscience). Cell confluence was calculated in 6 fields per well, in at least five replicates per treatment, in three independent experiments.

### Cell-by-Cell Analysis for detection of apoptosis

Apoptotic HAL and HuAST cells were detected using NucLight Rapid Red Reagent (Cat 4717) and Incucyte® Caspase-3/7 Green Dye (Cat 4440). Cells were treated with Vehicle (DMSO), T-DM1(1 μg/ml), TPM (100 μM) or T-DM1+TPM 2 hours before radiation (± 8Gy), using the RS2000 X-Ray irradiator. Plates were imaged for 3 days in a cell-by-cell module of Incucyte S3 (Essen BioScience). Analysis of apoptotic cells (green cells with red nuclei) was performed using the Incucyte software. n=6 replicates in at least two independent experiments. Details in supplemental methods.

### Western Blot (WB)

When indicated, cells for western blot were irradiated with 0-8Gy using the RS2000 X-Ray irradiator. PVDF membrane was then incubated overnight at 4°C with primary antibodies (**Sup. table 1)**. Membranes were imaged using Li-COR Odissey CLx and analyzed using Image Studio™ Software V5.2. Details in supplemental methods.

### Animal experiments

All animal experiments were approved by the University of Colorado Institutional Animal Care and Use Committee and the DoD-ACURO. Whole brain radiation therapy (WBRT) in female mice was performed using the Precision X-Ray X-Rad 225Cx Micro IGRT and SmART Systems, using 4.8 to 5.8 Gy/min in a single dose. To model radiation-induced brain edema, immune-compromised <u>N</u>OD-<u>S</u>CID IL2R<u>γ</u>null (NSG) female mice received 10Gy, and immunocompetent C57black/6J (Jackson Labs) female received 10Gy or 35Gy^18,19^ depending on the experiment. Experimental BM were developed by intracardially injection (ic) of 250,000 JmT1BR3-GFP-luciferase cells or 50,000 E0771BR1-GFP-luciferase cells in the left ventricle in 14-18-weeks old female mice (n=8 to 14 per group) and metastatic burden measured via IVIS. All animal interventions were performed by investigators blinded to the experimental groups.

### Brain water content

Freshly collected right brain hemispheres were put on a pre-weighed aluminum foil sheet, weighted (Melter Toledo, ME54E accuracy of 0.0001 g), dried at 99°C and weighed after 24 hours. The percentage of brain water content was determined as: (%) water content= 100×(wet weight-dry weight)/wet weight^20^.

### Electron microscopy

For astrocytic endfeet analysis, mice were anesthetized with isoflurane and intracardially perfused with PBS, followed by 2.5% glutaraldehyde, 2% PFA in 0.1 M Na-cacodylate pH 7.4. Brains were kept at 4°C in fixative solution and processed at the Electron Microscopy Center SOM (Details in supplemental methods). Images were acquired on a Tecnai T12 transmission electron microscope (ThermoFisher) equipped with a LaB6 source at 80kV using an XR80 (8Mpix) camera (AMT). Ten vessels/sample were imaged, and tight junctions from each vessel were imagined separately at higher magnification. Details in supplemental methods.

### Image Acquisition and Analysis

Microvessel images were collected using a 3i Marianas Spinning Disk Confocal microscope attached to a Zeiss Axio Observer Z1 with Yokogawa CSU-X1 microscope, with a 63x Oil objective. Maximum Z projections were made using SlideBook 6 software. For immunofluorescence (IF) analysis, images were collected using a Nikon Eclipse Ti-S inverted microscope using a Nikon CFI Plan Apo 20X/0.75 DIC M/N2 /0.17 WD 1.0 Objective MRD00205.

Image resolution was 1280 x 1024 px. Pixel size 0.32 μm/px. Region of interest were manually delineated using the drawing tool in Fiji software and mean Intensity and area of interest were calculated using the measure tool in Fiji software^21^. Details in supplemental methods.

### Transepithelial Electrical Resistance (TEER)

TEER in monolayer of HAL and HuAST plated on cell culture inserts was measured using a voltohmmeter (EVOM2, World Precision Instruments) with a chopstick electrode (STX3, World Precision Instruments). Measures were made before radiation and pre-treatments, and at the indicated times following radiation. The unit area resistance (Ωcm^2^) was obtained by multiplying the Ω reading by the surface area of the filter membrane (0.3 cm^2^). The TEER from empty inserts with media alone was subtracted from the TEER of cultures (values between ~160-200 Ω). Details in supplemental methods.

### Statistical Analysis

Specific statistical tests are listed in each figure legend, along with the number of samples assessed. Data statistical analysis and graphs were completed using GraphPad Prism software version 9.5.0. Randomization was used for all mice studies. All *in vitro* experiments were repeated at least three times unless indicated otherwise. Data was analyzed using unpaired t test, Mann-Whitney test, and two-way analysis of variance (ANOVA) with multiple comparison test as appropriate. A P value ≤0.05 was considered significant.

## RESULTS

### Radiation induced astrocytic swelling via AQP4 upregulation is an early step in brain edema

Consistent with our previous report^15^; immortalized human astrocytes (HAL and THV) and primary human astrocytes (HuAST) express baseline level of AQP4 (**Fig. 1A**) which transiently increases within the first 24 hours following radiation (8Gy, ***P=0.0325***) (**Fig. 1B, Sup. 1A**). To determine whether radiation-induced AQP4 upregulation resulted in water intake and astrocyte swelling, we measured the cell area of HuAST 24 hours post-radiation. A dose of 8Gy increased the astrocyte area by 4.8 fold compared with non-irradiated cells (***P<0.0001***) (**Fig. 1C**). Since radiation induces senescence in astrocytes and senescent cells are hypertrophic^22^, we assessed whether radiation-induced astrocytic swelling was due to senescence. Radiation increased the percentage of senescent HuAST (SA-β-Gal+) 48 hours after a single-8Gy dose (***P=0.002***) (**Fig. 1D**, **1E-left**). However, swollen astrocytes following 8Gy radiation were non-senescent (SA-β-Gal^neg^) (***P<0.0001***) **(Fig. 1E-right).** These results suggest that radiation-induced astrocytic swelling is not due to radiation-induced senescence.

**Figure 1:**
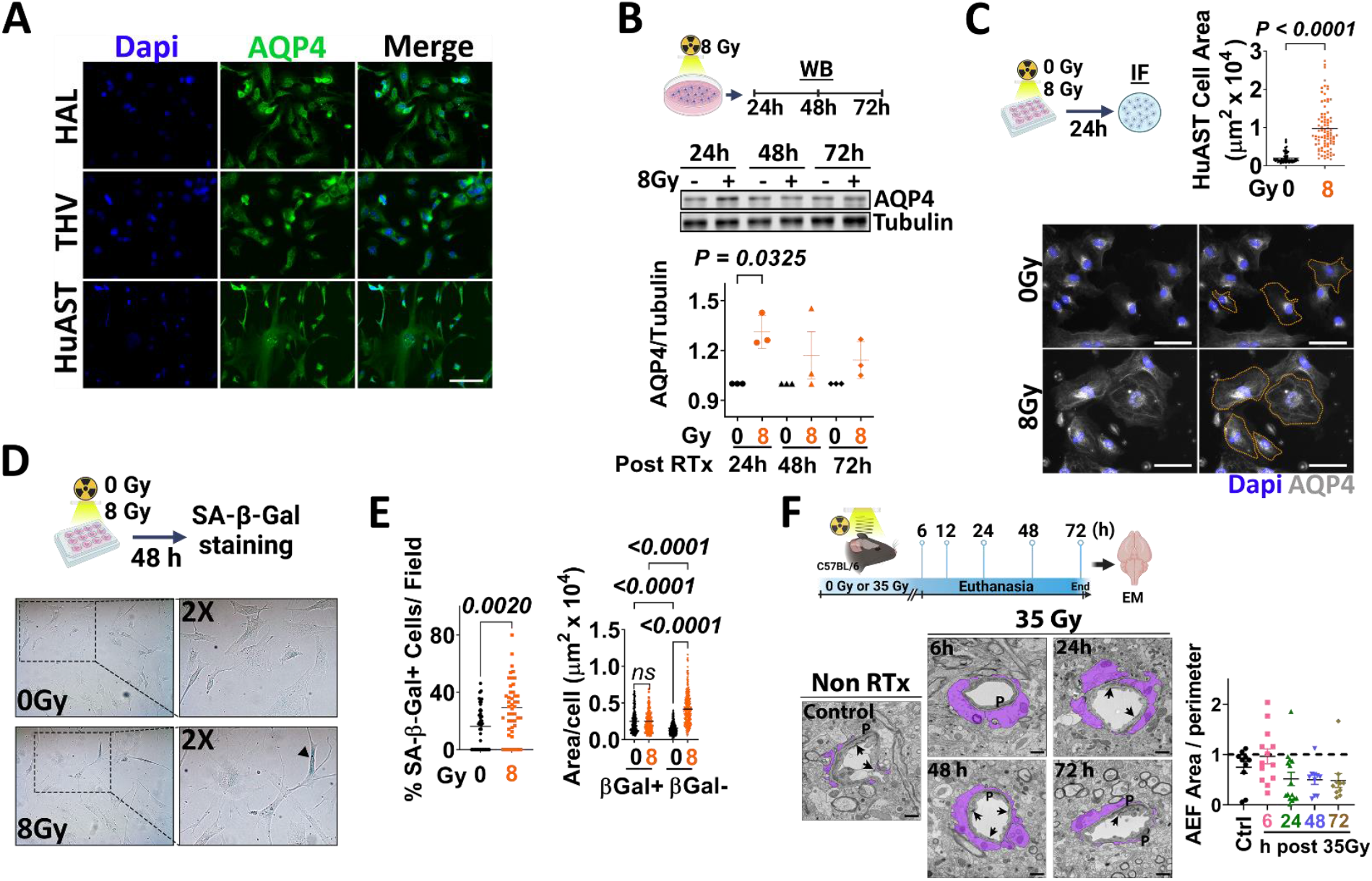
Radiation induces astrocytic swelling *in vitro* and *in vivo.* **(A)** AQP4 (green) expression in human astrocytes. Nuclei (Dapi-blue). Scale 100μm. **(B)** WB shows AQP4 levels inHuAST ± 8Gy at the indicated times. Graph shows fold change AQP4/tubulin levels relative to non-irradiated cells (n=3). Paired T-test. **(C)** Astrocytic area in AQP4-stained HuAST (gray) ±8Gy 24h post-radiation. Orange line shows astrocyte area delimitation. Scale 100μm. Graph shows area of single cells (n=86). **(D)** SA-β-gal staining in HuAST treated ± 8Gy. Blackhead arrow shows SA-b-gal+ cells. **(E)** *Left:* Percentage of SA-β-Gal+ HuAST ± 8Gy at 48h, quantified in 45 fields/condition on 3 independent slides. *Right:* Individual cell area of βGal+ or βGal-HuAST ± 8Gy at 48h, quantified in bright field images from 3 coverslips per condition using imageJ, analyzed by Kruskal-Wallis test followed by Dunn’s post-hoc test. **(F)** C57Bl6 mice were irradiated with 35Gy WBRT and euthanized at the indicated times. Representative microphotographs of cortex brain microvessels showing perivascular endfeet astrocytes (purple), pericytes (P), tight junction (black arrows) at the indicated times. Scale 1μm. Graph shows AEF area normalized to the vessel perimeter.

To determine if radiation-induced astrocytic swelling occurs *in vivo,* female C57Bl6 mice were treated with a single 35Gy-WBRT and the astrocytic end feet (AEF) surrounding microvessels in the brain cortex was analyzed using electron microscopy (**Fig. 1F).** A larger fraction of AEF with enlarged area (>1 AEF/vessel perimeter) was observed 6 hours following radiation compared to non-irradiated mice (46.5% vs 20.0% enlarged AEF, respectively) **(Fig. 1F)**, without significant changes in tight junction BBB integrity (**Sup. 1B**), suggesting that radiation induces astrocytic swelling and transient upregulation of AQP4 without leading to acute disruption of the BBB within the first 24 hours following radiation.

### Topiramate protects human astrocytes from radiation-induced cell swelling *in vitro*

AQP4 is critical for survival of astrocytes *in vitro*^23^. Consistently, stable knockdown of AQP4 using shRNAs prevented radiation-induced astrocytic swelling in HuAST but resulted in cell death **(Sup. 2A-C**). Thus, we used a pharmacologic approach to block AQP4 function and hypothesized that this will prevent radiation-induced cytotoxic swelling. We used a selective AQP4 inhibitor, TGN-020^24^, or an FDA-approved anti-epileptic drug Topiramate (TPM)^25^, previously reported to function as an AQP4 inhibitor^16^. Neither TGN-020 nor TPM induced cell death in THV astrocytes at 10-100 μM (**Sup. 2D**). Pre-treatment with TGN-020 slightly reduced the fraction of radiation-induced swollen astrocytes, but this effect was not statistically significant at the doses tested. By contrast, a 2 hours pre-treatment with 100 μM of TPM was effective in protecting THV from radiation-induced swelling (***P<0.0001***) (**Fig. 2A**). Therefore, we sought to determine if TPM could prevent astrocytic swelling in HuAST alone or in combination with cytotoxic therapies known to promote radiation-induced edema.

**Figure 2:**
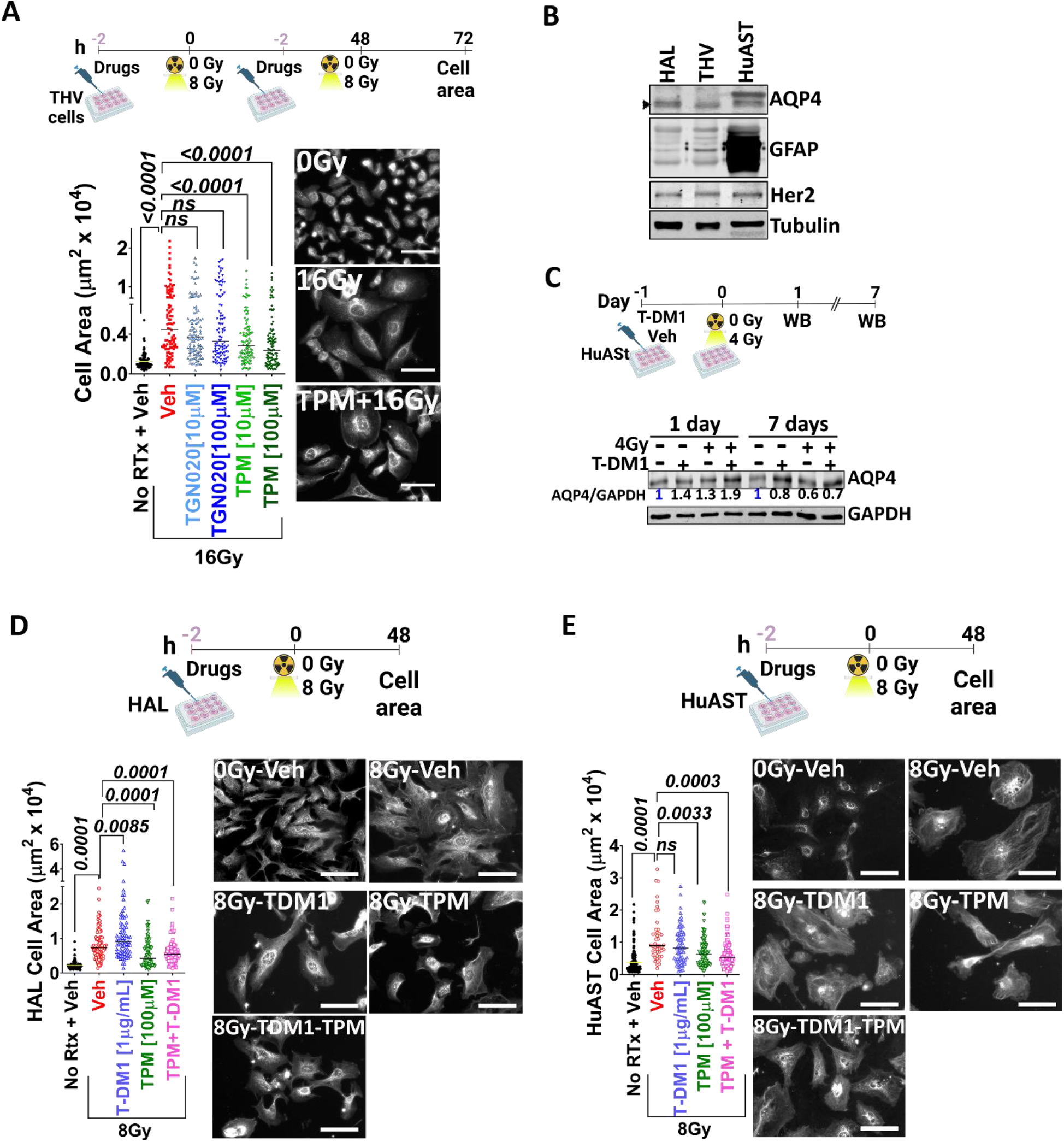
Topiramate prevents radiation-induced cell swelling in human astrocytes *in vitro.* **(A)** THV cells were pre-treated with Veh (DMSO), TGN020 (10-100μM) or TPM (10-100μM) 2h prior to ± 8Gy. Graph shows THV cell area 24h after 2 dose of 8Gy. IF images of THV stained with AQP4 (gray). **(B)** WB shows AQP4, GFAP and Her2 expression in human astrocytes. Tubulin is loading control**. (C)** HuAST were pre-treated with vehicle or T-DM1 [1μg/ml] 2h before ± 4 Gy. WB shows AQP4 levels 1 and 7 days post-radiation. GAPDH is loading control. Numbers are AQP4/GAPDH relative to untreated cells. **(D)** HAL cells were pre-treated with Veh (DMSO), T-DM1 (1μg/ml), TPM (100μM), or TPM+T-DM1, 2h prior to 8Gy. Graphs show cell area in μm^2^ 48h post-radiation. Representative images of AQP4 stained HAL cells (gray). **(E)** Cell size 48h post-irradiation in Hu-AST treated as in D. For all images, scale bar 100μm. Data were analyzed with Kruskal Wallis followed by post-hoc corrections (A) or Mann-Whitney test for paired comparisons (D,E).

We previously showed that the Her2-targeting agent T-DM1 increased radiation-induced astrocytic swelling in immortalized human astrocytes, which express normal levels of Her2 and are an unintended target of T-DM1^15^. Accordingly, T-DM1 increased AQP4 within 24 hours post-radiation in Her2+ HAL and HuAST, returning to normal levels 7 days post-radiation (**Fig. 2B-C**). To test whether TPM could prevent this transient dysfunction of AQP4, HAL and HuAST were pretreated with 100 μM of TPM for 2 hours before radiation and T-DM1 treatments (**Fig. 2D-E**). TPM prevented astrocytic swelling induced by radiation alone or in combination with T-DM1 in both HAL and HuAST cells, ***P<0.0001*** in HAL (**Fig. 2D**), and ***P=0.0033*** in HuAST (**Fig. 2E**).

To determine if preventing astrocytic swelling could also preserve astrocytic function in the BBB alone or in combination with T-DM1, we measured TEER of normal and irradiated astrocytic monolayers *in vitro* (**Fig. 3A**). Radiation changed TEER in a biphasic fashion; increasing TEER in the first 20 min (***P=0.0313*** in HAL and ***P<0.0001*** in HuAST) followed by a significant decrease after 24 hours (***P=0.0152***) (**Fig. 3A**). Therefore, we tested whether pre-treatment with TPM could prevent the loss of TEER 24 hours following radiation alone or in combination with T-DM1 (**Fig. 3B**). T-DM1 and radiation significantly decreased TEER compared to non-irradiated cells (HAL, ***P=0.0054***) and (HuAST, ***P<0.0001***) (**Fig. 3B**), and TPM pre-treatment restored TEER in T-DM1/Radiation-treated cells by 73% in HAL cells (***P=0.0459***) and 48% in HuAST (***P=0.0122***) as compared to T-DM1/Radiation-treated cells (**Fig. 3B**). Since radiation and T-DM1 can induce astrocytic cell death by mechanisms other than AQP4 dysregulation including direct DNA damage (**Sup. 3A**), we measured the impact of TPM in radiation and T-DM1 induced cell death. Using a live-imaging cell-by-cell detection of apoptosis **(Fig. 3C)**, T-DM1 and radiation alone and in combination significantly induced apoptosis by 72 hours compared to untreated cells in both HAL and HuAST (**Fig. 3D**). TPM showed a modest reduction in apoptosis in HAL cells and no effects in T-DM1/Radiation-induced apoptosis of HuAST. Consistent with these results, TPM did not affect radiation-induced double-strand DNA-damage as measured by pH2A.X activation, (**Sup. 3A**), caspase-3 activation (**Sup. 3B**) or activation of radiation-induced survival pathways (**Sup. 3C**). Thus, TPM can prevent astrocytic swelling and partially protects the physiological properties of astrocytes but cannot reverse radiation-induced cell death.

**Figure 3:**
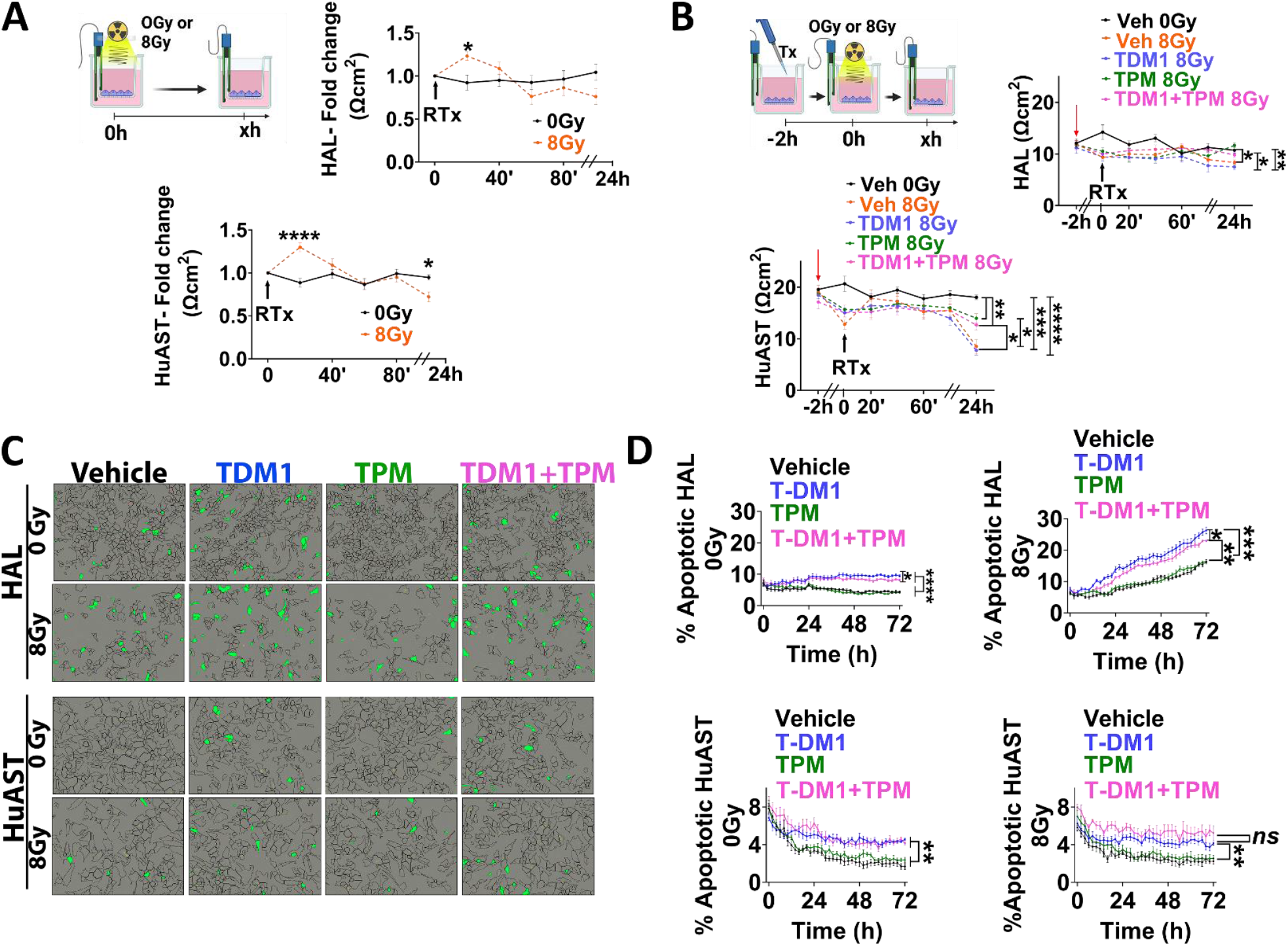
Topimarate restores radiation and TDM-1-induced loss of TEER but does not prevent astrocytic cell death *in vitro.* **(A)** Dynamics of TEER changes in monolayer of astrocytes grown on transwell inserts and irradiated with 8Gy. Graphs show TEER expressed as a fold change of Ωcm^2^ relative to time zero in HAL and HuAST. **(B)** TEER in a monolayer of astrocytes pre-treated with Veh (DMSO), T-DM1 (1μg/ml), TPM (100μM), or (TPM+T-DM1) 2h prior to 8Gy. Graphs show mean TEER in Ωcm2 (n=9 transwell/treatment). Lines are mean± SEM. **(C)** HAL or HuAST were treated as B. Representative images from Apoptotic and Caspase 3/7+ cells (green mask) 72h after treatment. **(D)** Graph shows percentage of apoptotic cells over time. Mean ± SEM (n=6 wells). *\*P<0.05, **P<0.01 ***P<0.001, ****P<0.0001.* (two-way ANOVA-Geisser-Greenhouse correction followed by post-hoc test).

### Short-term pre-treatment with Topiramate prevents radiation-induced brain edema *in vivo*

Changes in brain water content are widely used to measure water channel dysfunction and brain edema in pre-clinical mouse models^11,26^. To determine whether TPM could prevent radiation-induced astrocytic swelling and brain edema *in vivo*, we first measured water content in brains of untreated C57BL6 mice, or mice that had been exposed 24 hours before, to a single 35Gy dose of WBRT, pretreated or not with TPM (**Fig. 4A**). While radiation increased AQP4 in AEF in brain microvessels (**Fig. 4B**), there were no changes in brain water content 24 hours following irradiation in vehicle or TPM-treated mice as compared to untreated mice (**Fig. 4C**), suggesting that radiation-induced early astrocytic swelling does not result in acute changes in brain water content in the absence of BM.

**Figure 4.**
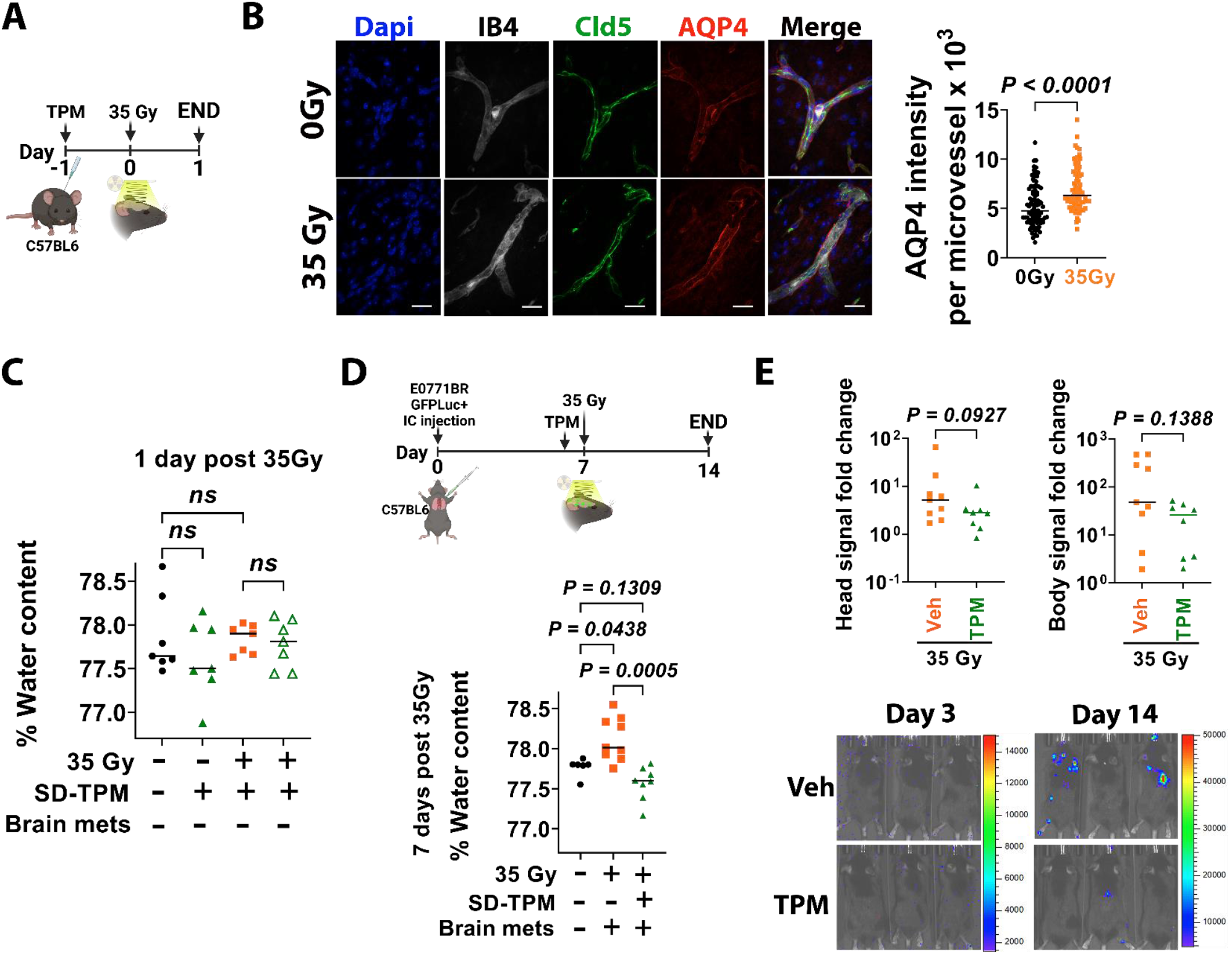
Pretreatment with topimarate prevent radiation induced brain water accumulation in a syngeneic model of BCBM. **(A)** C57BL6 mice were pre-treated with a single dose of vehicle or TPM [50mg/Kg] 24h prior to receiving or not 35Gy WBRT, and euthanized 24h later (n=10). **(B)** Confocal z-stack maximum projection of end-feet astrocytes AQP4+ (red) in IB4+ (Isolectin B4, gray) and Cld5+ (Claudin-5, green) microvessels. Dapi stains nuclei (Blue). Scale bar 20μm. Graph shows mean AQP4 intensity per brain microvessel (n=100 from three mice/treatment). Two-tail Mann Whitney test. **(C)** Percentage brain water content per mouse (n=6 per treatment) in mice from A. One way ANOVA followed by Sidak’s post-hoc test. **(D)** C57BL6 female mice were injected intracardially with E0771BR-GFP-Luciferase cells and metastasis allowed to grow for 7 days. Mice were then randomized to receive a single dose of TPM [50mg/Kg] or vehicle 4h prior to receiving 35Gy WBRT and euthanized 7 days later. Graph shows the percentage of brain water content per mouse (n=8,9 mice/group). Non-irradiated naïve C57BL6 mice (n=6) were included as baseline water content. Data were analyzed using Kruskal-Wallis followed by two-stage linear step-up procedure of Benjamini, Krieger and Yekutieli multiple comparisons test. **(E)** Fold change IVIS signal per mouse in head (left) and body (right). Mann-Whitney test. Representative images of IVIS signal 3 and 14 days after cell injection.

We next determined whether TPM could prevent radiation-induced edema in a pre-clinical model of BM. For this, breast cancer cells syngeneic to C57Bl6 mice (E0711-GFP-Luciferase) were injected intracardially in 8-12-week-old female mice, and BM were allowed to form for 7 days. Mice were then randomized to receive vehicle or TPM 4 hours before a single 35Gy WBRT, and brain water content was quantified 7 days later (**Fig. 4D**). In this model of radiation-induced edema in BM-carrying mice, TPM significantly reduced brain water content compared to vehicle-treated mice (**Fig. 4D**). Importantly, TPM prior to radiation did not affect brain metastatic burden (**Fig. 4E**), suggesting that short-term TPM is sufficient to prevent radiation-induced brain edema and does not increase BM burden in this model.

### Topiramate does not alter the response of cancer cells to radiation or T-DM1 treatment *in vitro*

While E0771 cells were not responsive to short-term TPM *in vivo*, AQP4 has been shown to play various roles in breast cancer cells^27,28^, and interventions targeting AQP4 have the potential to impact tumor burden. IHC staining in a cohort of breast cancer BM showed that AQP4 is heterogeneously expressed in BM (percentage positive tumor area ranging from 1.6% to 91.1%, median 10.5%, (**Fig. 5A, B**). AQP4 was also expressed in 5 of 7 metastatic breast cancer cell lines from all breast cancer subtypes, as well as in murine mammary cancer models (**Fig. 5C**). Moreover, radiation modified AQP4 expression in a cell line-dependent manner (i.e radiation increased AQP4 expression in MCF7 and E0771 cells but reduced its expression in 4T1BR5 cells (**Fig. 5C**).

**Figure 5:**
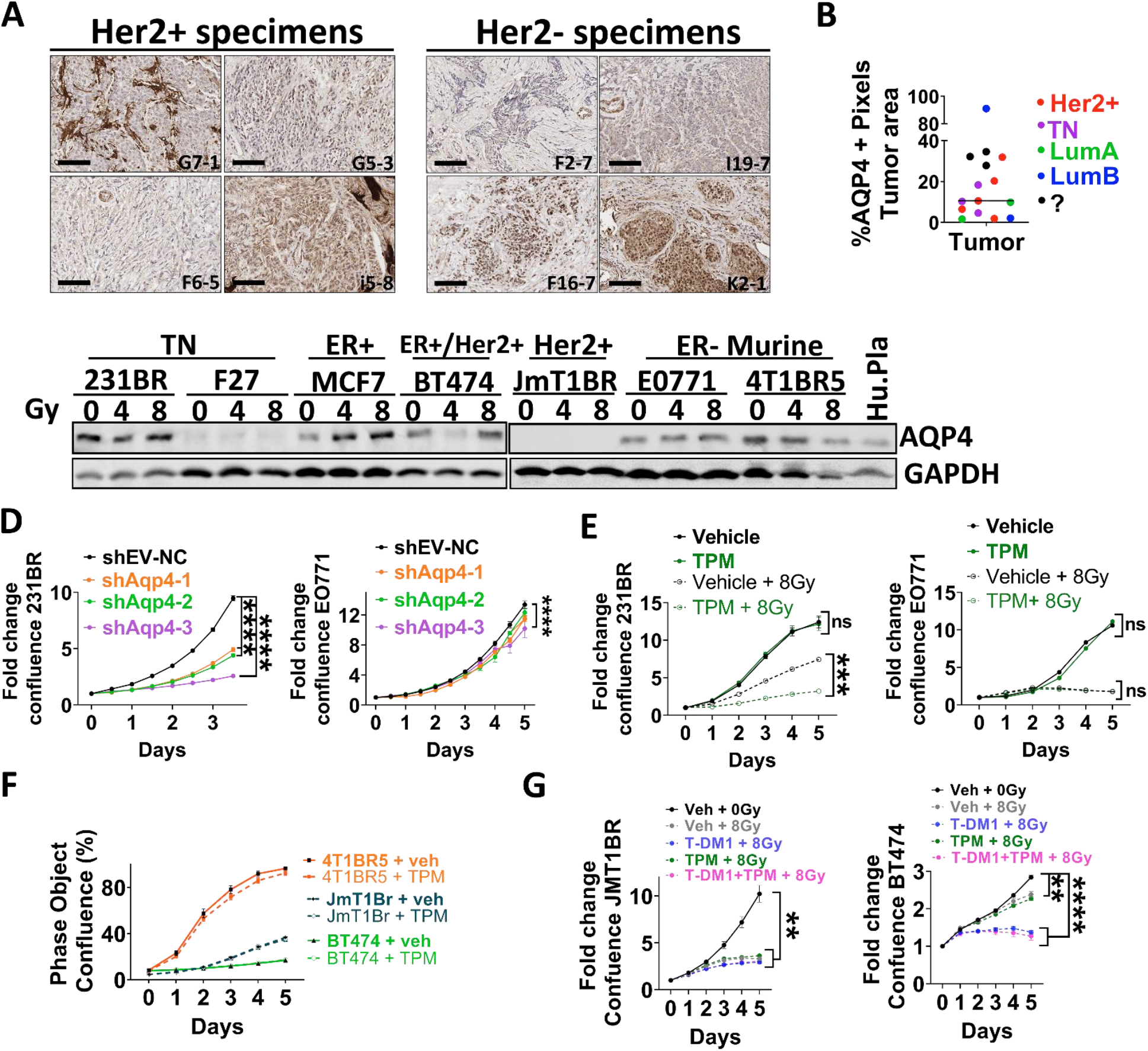
AQP4 is expressed in breast cancer brain metastasis. **(A)** AQP4 IHC staining in Her2^+^ and Her2^-^ BCBM clinical samples. **(B)** Percentage of positive pixels in tumor area in a cohort of 14 BCBM patients from the indicated subtypes. **(C)** AQP4 WB in human and murine breast cancer cells treated with 0, 4 and 8Gy. **(D)** Fold change confluence of 231BR or E0771 cells infected with shRNA shEV or three shRNAs targeting human (231BR) or murine (E0711) AQP4, respectively. **(E)** Fold change in confluence of 231 and E0771 cells treated with vehicle (DMSO) or TPM [100μMl] 2h prior ± 8Gy. **(F)** Percentage confluence of indicated breast cancer cells treated with Vehicle (DMSO) or TPM [100μM]. **(G)** Her2+ JmT1BR or BT474 cells treated with vehicle (DMSO), T-DM1 [1 μg/ml], TPM [100 μM], or TPM+T-DM1, 2h prior to 8Gy. For all graphs, line is mean ± SEM. Data were analyzed using Two-way ANOVA with the Geisser-Greenhouse correction followed by Tukey’s multiple comparisons test.

To test whether targeting AQP4 could directly influence cancer cell proliferation, AQP4 was knocked down in 231BR and E0771BR cells, using three different lentiviral shRNAs targeting human or murine AQP4, respectively. AQP4 knockdown in the human 231BR cells decreased their proliferation significantly compared with control cells (***P<0.0001***), while AQP4 knockdown in the murine E0771 cells had negligible effects on cell proliferation (**Fig. 5D**). Consistently, TPM increased the sensitivity to radiation in 231BR, but had no effect on E0771 cells (**Fig 5E**). Moreover, TPM treatment did not impact proliferation of other breast cancer cell lines (**Fig. 5F**), nor it impacted sensitivity to T-DM1 and radiation in two Her2+ cell lines (**Fig. 5G**). Taking together, these results suggest that TPM does not have direct pro-tumorigenic effects, neither does it have a radioprotective effects in cancer cells that could impact its clinical translation in preventing radiation-induced brain edema.

### Long-term Topimarate decreased brain edema but increased tumor burden in syngeneic mouse models but not in immunocompromised models

Since TPM is an FDA-approved anti-epileptic drug used daily, we sought to determine whether a daily TPM administration could improve long-term control of radiation-induced brain edema in pre-clinical models of BM. We chose models that were not sensitive to TPM *in vitro* (JmT1Br3 and E0711) to be able to measure the effects of TPM on radiation-induced edema independent on tumor burden. First, we injected AQP4^neg^ human JmT1Br3 cells in severely immunocompromised NSG mice, allowed metastasis to form for 7 days, and then randomized mice to receiving or not a single 10Gy dose (maximum permissible dose in NSG mice) and a daily dose of TPM or vehicle for the following 14 days (**Fig. 6A**). Consistent with the results found using short-term use of TPM in the E0711/C57BL6 model, sustained TPM treatment reduced brain water content following irradiation (**Fig. 6A**) without changes in tumor burden in the brain or in systemic metastasis (**Fig. 6B**). Unexpectedly, sustained TPM treatment in C57Bl6 mice carrying E0711 BM reduced brain water content in irradiated mice (**Fig. 6C**) but led to a significant increase in metastatic tumor burden (**Fig. 6D**). Since TPM did not directly alter the growth of E0711GFP-Luc cells *in vitro* (**Fig. 5F**), and this pro-metastatic effect was not observed in the JmT1BR3/NSG model, we proposed that the immune environment was responsible for this pro-tumorigenic effect of long-term TPM. Consistent with this hypothesis, sustained TPM in NSG mice injected with E0711-GFP-Luciferase cells (**Fig. 6E**) resulted in decreased water content (**Fig. 6E**) but did not lead to increases in metastatic burden (**Fig. 6F**). Thus, long-term use of TPM in the context of metastatic breast cancer may impact metastatic burden through modulation of the immune microenvironment.

**Figure 6.**
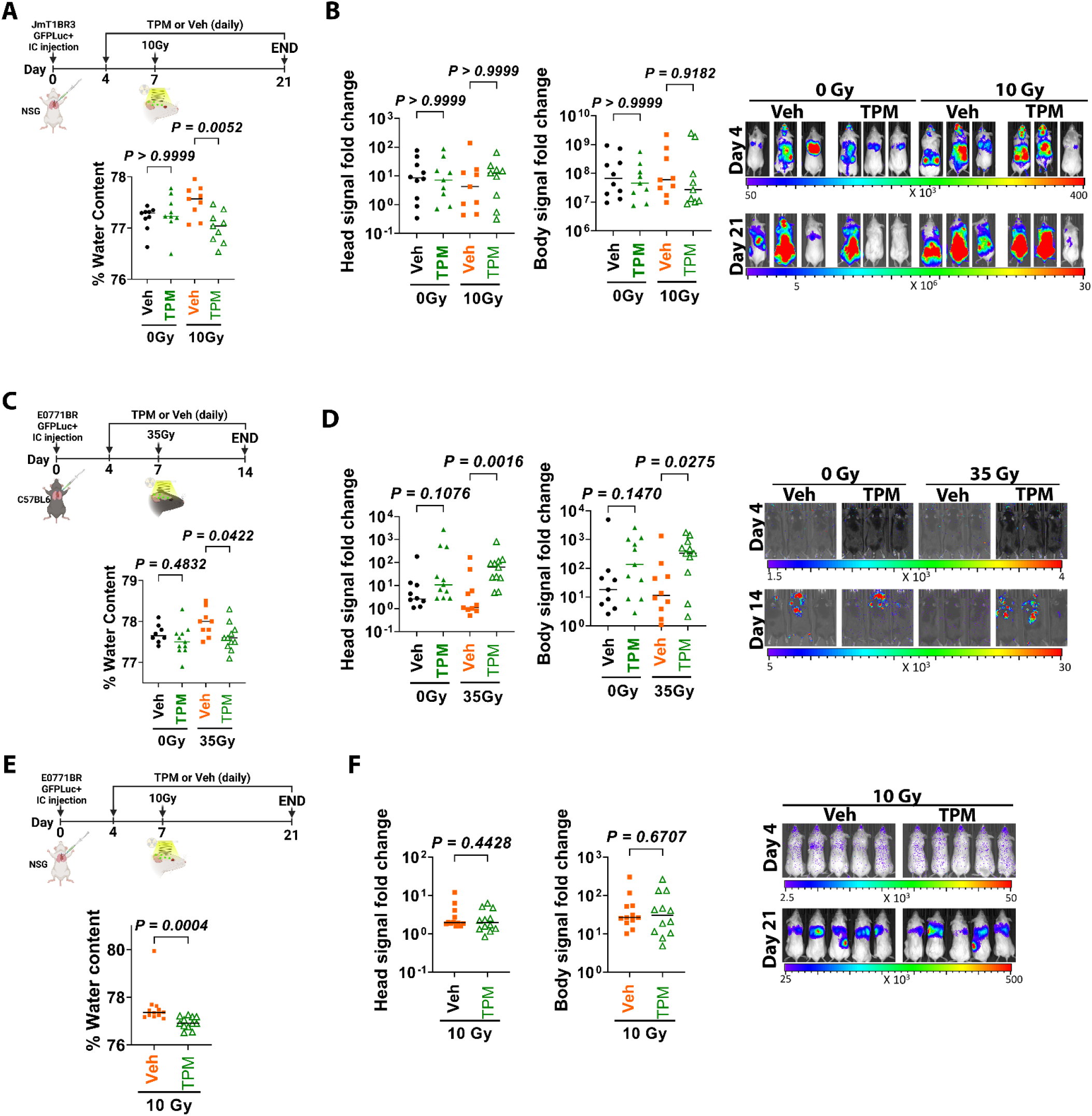
Long-term Topimarate decreased brain edema but increased tumor burden in syngeneic mouse models but not in immunocompromised models. **(A)** NSG mice were injected intracardially with JmT1BR3-GFP-Luciferase cells and 4 days later randomized to receive daily doses of vehicle or TPM [50mg/Kg]. Mice received or not 10Gy WBRT 7 days post-injection and were euthanized 14 days later. Graph shows percentage of brain water content per mouse (n=9 per group). **(B)** Fold change metastatic burden in the head (left) and body (right) measured via IVIS in mice from A. Panel shows representative IVIS images 4 and 21 days after cancer cell injections. **(C)** C57BL6 mice were injected intracardially with E0771BR-GFP-Luciferase cells and treated as in A, except that received 35Gy WBRT on day 7 and euthanized 7 days later. Graph shows percentage of brain water content per mouse (n=9 to 11/group). **(D)** Fold change metastatic burden in the head (left) and body (right) measured via IVIS in mice from C. Panel shows representative IVIS images 4 and 14 days after cell injections. **(E)** NSG mice were injected intracardially with E0771BR-GFP-Luciferase cells and treated as in A, except that all mice received 10Gy WBRT. Graph shows percentage of brain water content per mouse (n=12/group). **(F)** Fold change of metastatic burden in the head (left) and body (right) measured via IVIS in mice from E. Panel shows representative IVIS images 4 and 21 days after cancer cell injections. (A-D) Data was analyzed using Kruskal-Wallis followed by Dunn’s post-hoc test. (E-F) was analyzed using Mann-Whitney test.

## DISSCUSION

In this study, we describe a novel role for AQP4 and cytotoxic edema in BM and radiation-induced brain edema. While edema in the context of BM has been fully attributed to tumor-induced disruption of the BBB leading to vasogenic edema, our results suggest that cytotoxic edema affecting astrocytes also plays an important role in the progression of brain edema during BM standard of care.

Using *in vitro* models of human astrocytes and *in vivo* models of WBRT, we show that radiation induces astrocytic swelling and transient upregulation of AQP4, hallmarks of cytotoxic edema in other brain injuries^10,29–32^. We show that radiation-induced astrocytic swelling is not occurring in senescent cells; thus, increased astrocytic cell size can be attributed to disrupted water intake and not hypertrophy occurring during senescence^22,33^. Moreover, the fact that AQP4 blockage can decrease astrocytic swelling and restore TEER function *in vitro* but does not protect from cell death induced by radiation or cytotoxic drugs such as T-DM1, further suggest that acute cytotoxic edema and AQP4 dysfunction are not mechanistically linked to DNA damage or microtubule disruption, the main cytotoxic drivers of radiation and T-DM1, respectively.

In contrast to prior reports showing that 35Gy of WBRT on C57BL/6J mice induced brain water accumulation after 1 day of radiation^19^, we observed that a 35Gy dose lead to AQP4 upregulation *in vivo*, without leading to acute disruption of the BBB or changes in brain water content within the first 24 hours following radiation. Nonetheless, radiation doses between 10-35Gy induced a significant increase in brain water content in BM-carrying mice, and a single pre-treatment dose of TPM administered within 4 hours from radiation were able to block this effect, further supporting the hypothesis that acute cytotoxic edema plays a role in the pathobiology of radiation-induced edema and can be targeted in the context of BM.

Acute inhibition of AQP4 has been shown to decrease edema and infarct lesion volume in pre-clinical models of stroke^34^, however, the existing AQP4 specific inhibitors are far from clinical application. In this study we use TPM due to its previously reported role in blocking AQP4-dependent water flux^16^, and the fact that is an FDA-approved anti-epileptic drug with rapid translational potential. However, we recognize that TPM has additional functions and may decrease cytotoxic edema through additional mechanisms. Recent studies showed that AQP4 subcellular re-localization to the plasma membrane is required to drive cytotoxic edema in models of hypoxia-induced cell swelling^11^. AQP4 translocation to the membrane was shown to require an influx of Ca^2+^ ions to activate calmodulin, which in turn activates adenylyl cyclases leading to PKA activation and subsequent phosphorylation of AQP4, ultimately responsible for AQP4 translocation to the plasma membrane^11^. Since TPM targets calcium channels and TPM has been reported to decrease Ca^2+^ influx in neurons^35^, it is possible that TPM alters the intracellular concentration of Ca^2+^ also in astrocytes, thus, reducing AQP4 function at the plasma membrane. This potential mechanism for TPM in blocking AQP4 function could explain why TPM but not the AQP4 specific inhibitor TGN-020 which targets AQP4 channel blocker function, decreased radiation-induced astrocytic swelling *in vitro*.

Patients with BM are at significant risk for developing seizures^36^, and anti-epileptic drugs are prescribed to control this side effect. Thus, we tested whether daily TPM administration could improve long-term control of radiation-induced brain edema in pre-clinical models of BM. Unexpectedly, our results showed that long-term use of TPM in the context of BCBM increased systemic metastatic burden, raising concerns about the potential side effects of TPM and other anti-epileptic drugs targeting AQP4 in patients with systemic metastasis. While E0771 cells express AQP4, neither AQP4 KD or TPM altered E0711 cells proliferation *in vitro*, and TPM did not alter E0711 growth in NK and T-cell depleted NSG mice, suggesting that long-term TPM treatment promotes metastatic progression through modulation of immune cells. It has been shown that AQP4 is expressed on murine CD4+ and CD8+ T cells, and that blockage of AQP4 using a small molecule (AER-270) inhibits T cell proliferation *in vitro* and alters T-cell trafficking *in vivo* in a model of cardiac transplantation^37^. Thus, we propose long-term TPM or AQP4 inhibitors may suppress anti-tumoral T-cell function and thus indirectly facilitate metastatic tumor growth. Further studies are warranted to decipher the mechanisms whereby TPM and other anti-epileptic drugs targeting AQP4 could influence immune cells and tumor progression.

Expression of aquaporins in brain tumors and other malignancies has been shown to promote migration and proliferation^38^ and can alter the response of conventional anticancer therapies^27,39^; however, little is known about AQP4 function in BCBM. While AQP4 was expressed in several breast cancer cell lines, the majority were not susceptible to AQP4 blockage (either genetic or via TPM), suggesting that TPM or AQP4 blockage can be used to prevent cytotoxic edema without direct pro or anti-tumoral effects.

Finally, our studies have important clinical implications. First, current standard of care for brain edema involves post-edema use of bevacizumab and steroids targeting vasogenic edema. Our results suggest that patients with BM could find additional benefits from acute and temporary preventive treatment of radiation-induced cytotoxic edema using an already FDA-approved anti-epileptic drug. The proposed short use of TPM is similar to current clinical use of steroids in this setting. Lastly, our studies highlight the need for better understanding of the interactions between drugs commonly used in the care of patients with BM and their cancer-specific treatments.

## Supporting information

Supplementary Table 1 Antibodies

Supplementary Figures with Legends

## ACKNOWLEDGMENTS

We thank Dr. A. Van Bokhoven at the Biorepository Core Facility, and personnel at the University of Colorado Neurosurgery Nervous System Biorepository for providing de-identified human tissues. Dr. D. Yu for providing breast cancer cells BT474, and Dr. P. Steeg for providing 231BR and JmT1BR3 cells. The University of Colorado Cancer Center shared resources supported by NCI P30CA046934 and CTSA UL1TR001082 Center grants, the Advanced Light Microscopy Core at University of Colorado Anschutz Medical Campus supported by Rocky Mountain Neurological Disorders Core P30 NS048154 and by Diabetes Research Center P30 DK116073. Special thanks to Benjamin Van Court and Brooke Neupert in the Small-Animal Irradiator Shared Resource, Dr. Jennifer Bourne (†) and Dr. Anza Darehshouri in the Electron Microscopy Center at University of Colorado Anschutz Medical Campus.

