## Supplementary Table 1 Antibodies for "Short-term Topiramate treatment prevents radiation-induced cytotoxic edema in preclinical models of breast-cancer brain metastasis"

**Supplementary Table 1: List of antibodies**

| Target | Vendor | Cat# | Lot | Dilution/Use | RRID |
|---|---|---|---|---|---|
| AKT | Cell Signaling | 9272S | 25 | 1:1000/WB | AB_329827 |
| pAKT(S473) | Cell Signaling | 4060S | 25 | WB | AB_2315049 |
| AQP4 | Abcam | ab46182 | GR3204332-1 | 1:1000/WB | AB_955676 |
| AQP4 | Millipore | AB3594 | 3072342 | 1:1000/WB; | AB_91530 |
| Cleaved Caspase 3 (D175) | Cell Signaling | 9661 | 43 | IF | AB_2341188 |
| ERK | Cell Signaling | 9102 | 23 | WB | AB_330744 |
| pERK(T202/T204) | Cell Signaling | 9101 | 27 | WB | AB_331646 |
| GAPDH | cell signaling | 97166 | 5 | WB | AB_2756824 |
| GFAP | Thermo Fisher | 13-0300 | TA265137 | 1:1000/WB; 1:400/IF | AB_2532994 |
| pH2A.X(S139) | Cell Signaling | 80312 | 1 | IF | AB_2799949 |
| Her2 | Thermo Fisher | MA5-14509 | TK2671181B | WB | AB_10980124 |
| JNK | Cell Signaling | 9252 | 17 | WB | AB_2250373 |
| pJNK(T183/T185) | Cell Signaling | 9255 | 33 | WB | AB_2307321 |
| P38 | Cell Signaling | 8690 | 8 | WB | AB_10999090 |
| pP38(Y180/Y182) | Cell Signaling | 9216 | 27 | WB | AB_331296 |
| PARP | cell signaling | 9532 | 9 | WB | AB_659884 |
| PLCγ | SantaCruz | sc-81 | E1613 | WB | AB_632202 |
| pPLCγ(Y383) | Cell Signaling | 14008 | 3 | WB | AB_2728690 |
| Tubulin | Sigma | T5168 | 035M4878V | WB/IF | AB_477579 |
| VEGF | MIllipore | ABS82 | 3125474 | WB | AB_10806337 |
| Goat anti-Mouse Alexa Fluor 680 | Life Technology | A21058 | 2115694 | 1:10000/WB | AB_2535724 |
| Goat anti-Rabbit Alexa Fluor 680 | Life Technology | A21109 | 2260898 | 1:10000/WB | AB_2535758 |
| Goat anti-mouse Alexa Fluor 800 | LICOR | 926-32210 | C70301-02 | 1:10000/WB | AB_621842 |
