## Supplementary Figures with Legends for "Short-term Topiramate treatment prevents radiation-induced cytotoxic edema in preclinical models of breast-cancer brain metastasis"

**SUPPLEMENTARY FIGURE 1**

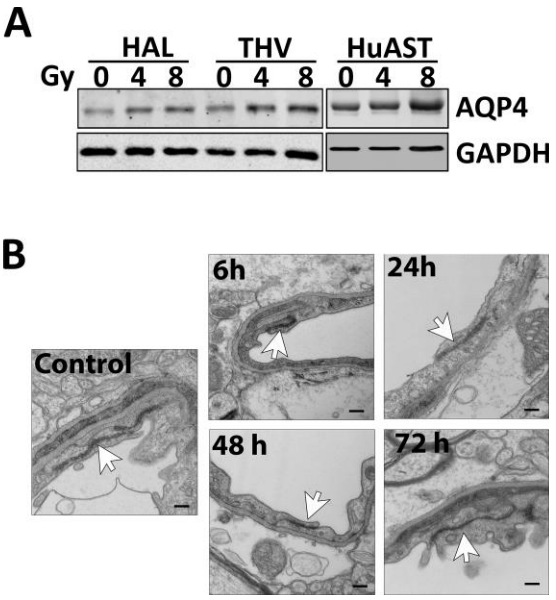


**Sup. 1: (A)** WB of AQP4 in human astrocytes treated with increased dose of radiation. GAPDH was used as loading control. **(B)** Representative microphotographs of cortex brain microvessels showing tight junctions (black arrows) at the indicated times.

**SUPPLEMENTARY FIGURE 2**

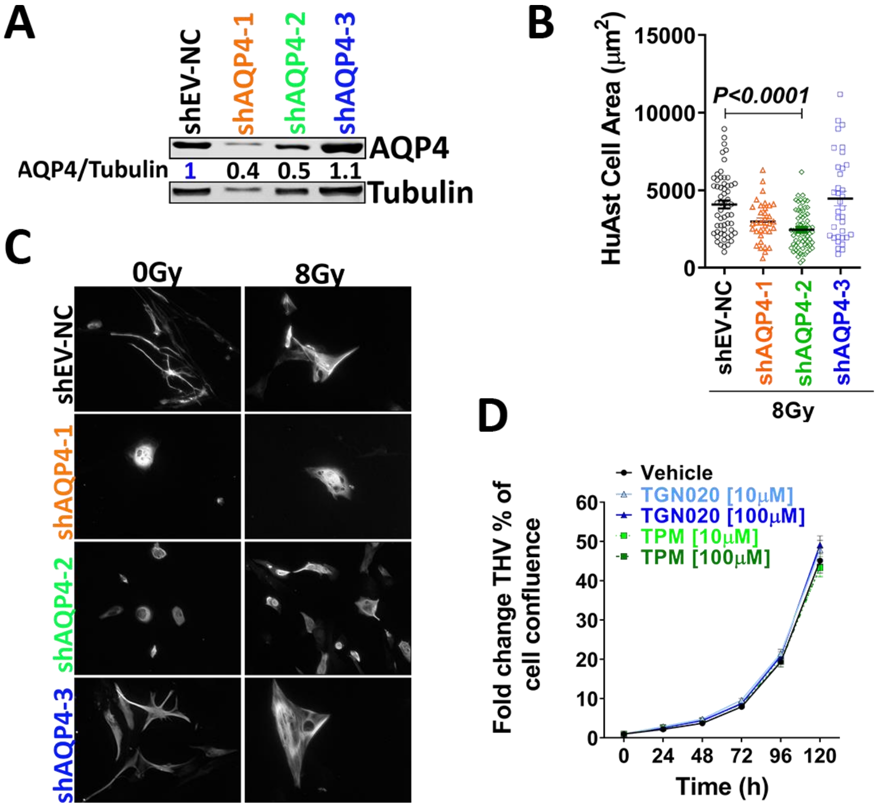


**Sup. 2: (A)** WB shows AQP4 levels in HuAST transduced with shRNA lentiviral vectors targeting human-AQP4 (shAQP4-1,-2 and -3) or a non-targeting control (shEV-NC) 48 h after infection, transduced positive cells were selected for 15 days with 1 µg/ml Puromycin. Numbers indicate the ratio of AQP4/α-tubulin relative to shEV-NC. **(B)** Graph shows cell area of single cells in A, 48 h after 8Gy. **(C)** Representative IF image of transduced shAQP4 HuAST stained for GFAP (gray). **(D)** Data shows mean fold change ± SEM of the percentage phase object confluence for THV cells treated TGN020 (10-100 µM), TPM (10-100 µM), or Vehicle (DMSO), relative to at time zero. Until 120 h.

SUPPLEMENTARY FIGURE 3

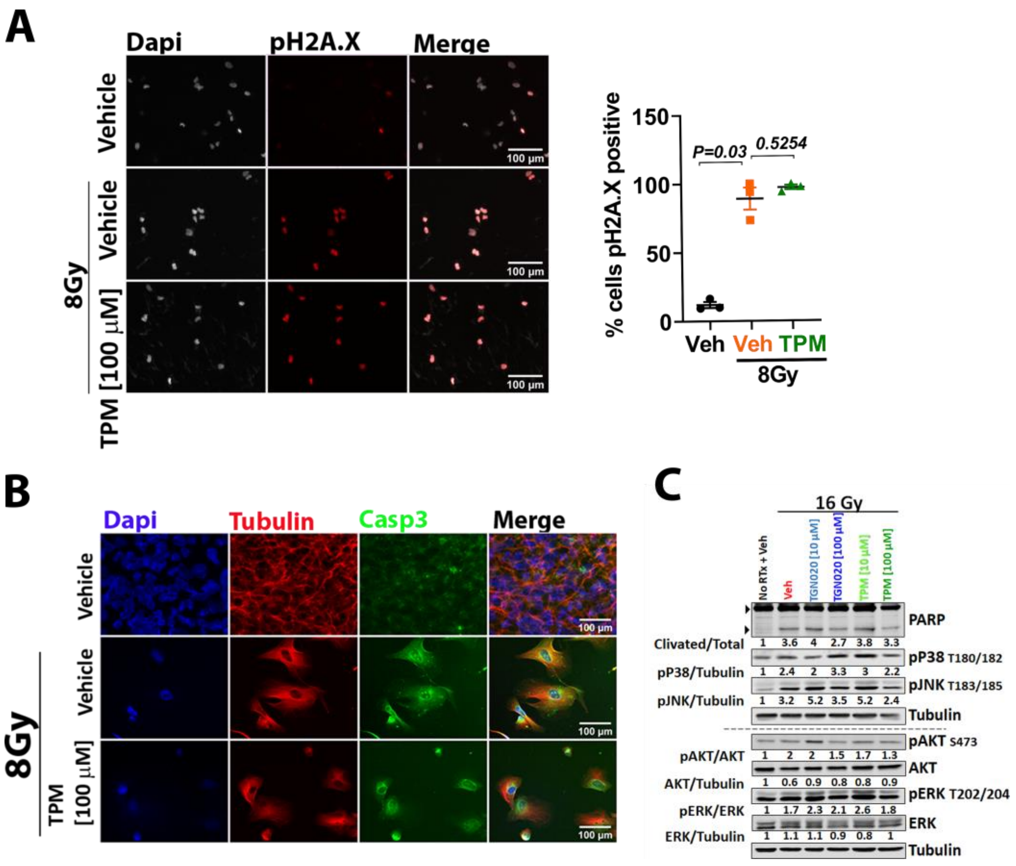


**Sup. 3: (A)** IF of pH2A.x a DNA double-strand breaks marker in THV cells pretreated for 2 h with TPM [100 µM] before 8Gy. Graph shows media of percentage of nuclei positive for pH2A.X. at least 100 nuclei were analyzed by condition. Kruskal-Wallis one-way ANOVA and Dunn post-hoc analysis. Bar scale 100 µm. **(B)** IF of Caspase 3 and tubulin in THV cells. Cells were plated in coverslips and pretreated for 2 h with indicated doses of Vehicle or TPM before a single dose of radiation (8Gy). Cells were fixed 48 h after radiation treatment. **(C)** Representative WB of MAPK proteins from THV cells treated AQP4 inhibitors 2 h before radiation. Tubulin was used as a loading control. Numbers indicate the ratio of protein/Tubulin to non-irradiated- non-treated cells.

### SUPPLEMENTARY FIGURE 4

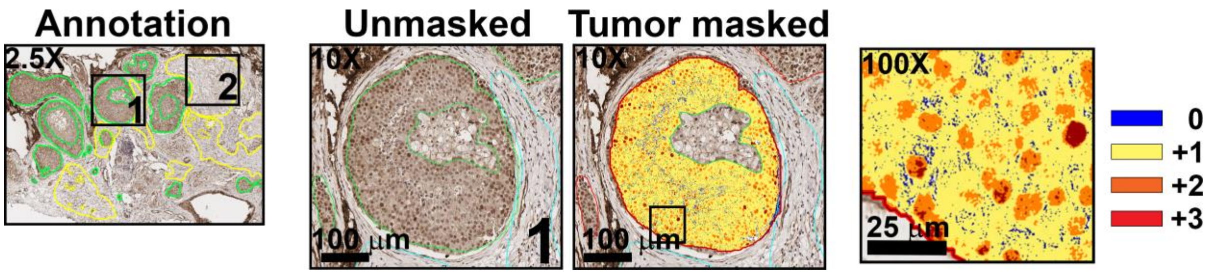


**Sup. 4:** Aperio software analysis for quantification of AQP4 in clinical samples**.** Entire histological sections were imaged with Aperio ScanCope T3 scanner at 0.25 μm/pixel. Sample areas for quantification were annotated using Aperio analysis tools and a minimum of 4 $mm^2$ tumor area (5000-20.000 tumor cells) were annotated from each sample. Stroma and necrotic areas were excluded from analysis. Algorithms developed in the Pathology Department at University of Colorado for AQP4 staining in clinical samples were used to assess staining of AQP4. Clinical samples with scores of 2+ (moderate Intensity, orange masks) and 3+ (strong intensity, red masks) were considered positive.
