## Supplementary material for "Short-term Topiramate treatment prevents radiation-induced cytotoxic edema in preclinical models of breast-cancer brain metastasis": Methods supplements

**Cell culture conditions:** HuAST and HAL astrocytes were maintained in astrocyte media (AM media ScienceCell #1801) supplemented with astrocyte-growth factor (AGS ScienceCell #1852), 2% FBS (ScienceCell #0010), and 100 units/ml of Penicillin-Streptomycin (P/S) (complete AM media). THV astrocytes were maintained in high glucose Dulbecco's Modified Eagle Medium (Corning 10-013-CV) supplemented with 10% FBS and 100 units/ml P/S. BT474 were cultured in F12 media with 10% FBS and 100 units/ml. MCF-7 cells were maintained in MEM plus 10% FBS, 2 mM of L-Glutamine, and 1 mM of Sodium Pyruvate. JMT1BR3 and 231BR cells were cultured in high glucose DMEM supplemented with 10% FBS and 100 units/ml P/S. F2-7 cells were maintained as described in Contreras-Zárate et. al[19]. 4T1BR5, a gift from Dr. Suyun Huang (MD Anderson Cancer Center, TX, USA) was cultured in high glucose DMEM supplemented 5 % FBS. E0771BR cells were maintained in DMEM 5% FBS and 100 units/ml P/S. 4T1BR5 was cultured in high glucose DMEM supplemented with 5 % FBS and 100 units/ml P/S. All experiments were performed on cells within 20 passages from thawing; cultures were tested to be free of mycoplasma, and cell identity was verified by STR analysis before freezing initial stocks

**Electron microscopy:** Samples were rinsed in 0.1M Na-cacodylate buffer and postfixed in 1% osmium tetroxide and 1.5% Potassium Ferrocyanide in 0.1M sodium cacodylate buffer for 15 min. They were rinsed again in 0.1M sodium cacodylate buffer and post-fixed in 1% osmium tetroxide in 0.1M Na-cacodylate buffer for one and a half hours at room temperature. After five rinses in 0.1M Na-cacodylate buffer and two rinses in water, samples were en bloc stained with 2% uranyl acetate in water overnight. Next, samples were dehydrated with increasing concentrations of ethanol, transitioned into resin with propylene oxide, infiltrated with EMbed-812 resin and polymerized in a 60 °C oven overnight. Blocks were sectioned with a diamond knife (Diatome) on a Reichert ultramicrotome and collected onto copper grids, post-stained with 2% aqueous uranyl acetate and lead citrate.

**Western blot:** Cells for western blot (WB) studies were plated (250,000 – 500,000 cells/60 mm dish; 100,000 cells/ 35 mm dish). Cell pellets were lysed at 4 °C for 5 min using 2X RIPA lysis buffer containing 2X protease inhibitor cocktail (Roche, 04693159002) and 2X phosphatase inhibitor cocktail (Roche, 04906837001), followed by 4 X 1-s sonication pulses at 20% amplitude. Protein concentration was determined using BioRad DC protein assay kit II (#5000112). Between 20 and 40 µg of protein samples were run in a polyacrylamide gel at 100 V in 10% SDS gels or 4-15% precast gels (Biorad Cat #4561086). After electrophoresis, proteins were transferred to an Immobilon-FL PVDF membrane 0.45 µm (Millipore # IPFL00010) for protein > 40 KDa or PVDF 0.2 µm (Amersham Hybond #10600022) and incubated in blocking buffer (3% BSA in TTBS) for 60 min at room temperature.

**Image Acquisition and Analysis:** 20 µm mouse brain sections were used for 3D brain microvessel analysis. Whole microvessels were imaged at 0.3 µm/step with 1x zoom and 0.169 µm/pixel XY size averaged 4 times. All acquisitions were made with 0 gain, 1x1 binning, and 150 camera intensification. For IF images, resolution was 1280 x 1024 px. Pixel size 0.32 µm/px. Digital images were exported as TIFF files to Adobe Photoshop. All manipulations and adjustments were performed identically and in parallel for each experiment.

**Transepithelial Electrical Resistance (TEER):** 100,000 HAL or HuAST were plated in 700 µl of complete AM media on cell culture inserts (0.4 µm pore size, Corning #353095) in 24 well plates containing 1 ml of complete AM media/well. Experiments were performed four days after plating when astrocytes formed a 100% confluent monolayer. Due to hand movement or temperature changes, the chopstick electrode was fixed to a support and plates were on a heating pad to avoid discrepancies.

Resistance in cells/inserts not irradiated was determined simultaneously with irradiated cells/inserts to control for the integrity of the monolayer.

**Cell-by-Cell Analysis for detection of apoptosis:** 1,500 HAL or HuAST cells/well were plated in complete AM media on a 96 well plate and treated for 2 hours before radiation (8Gy) with media containing NucLight Rapid Red Reagent for nuclear labeling (1:500) and Incucyte® Caspase-3/7 Green Dye for Apoptosis (5 μM), in combination with Vehicle (DMSO), T-DM1(1 μg/ml), TPM (100 μM) or T-DM1+TPM.
